# Prenatal Alcohol and THC E-Cigarette Exposure Effects on Motor Development

**DOI:** 10.1101/2021.05.26.445823

**Authors:** Kristen R. Breit, Cristina Rodriguez, Annie Lei, Samirah Hussain, Jennifer D. Thomas

## Abstract

It has been well established that prenatal alcohol exposure can lead to a wide range of neurological and behavioral deficits, including alterations in motor domains. However, much less is known about the effects of prenatal cannabis exposure on motor development, despite the fact that cannabis is the most commonly consumed illicit drug among women. Cannabis use among pregnant women has become increasingly popular given the wide-spread perception that consumption is safe during pregnancy. Moreover, alcohol and cannabis are commonly used together, even among pregnant women. Yet, few studies have explored the potential consequences of combined prenatal exposure on behavioral domains. Using our previously established model, pregnant Sprague-Dawley rats were exposed to vaporized alcohol, THC via e-cigarettes, the combination, or a vehicle from gestational days 5-20. Following birth, offspring were tested on early sensorimotor development, adolescent motor coordination, and adolescent activity levels. Prenatal e-cigarette THC exposure delayed sensorimotor development early in life and impaired motor coordination later in adolescence. However, combined prenatal alcohol and THC exposure produced hyperactivity among male offspring. These data suggest that prenatal cannabis exposure may lead to impaired motor skills throughout early development, and that combined exposure with alcohol during gestation may also lead to hyperactivity in adolescence. These findings have important implications for pregnant women and public policy.

## 1.1 Introduction

It is well known that prenatal alcohol exposure can lead to a wide range of physical and behavioral alterations, known as fetal alcohol spectrum disorders (FASD). The prevalence rates of FASD are estimated to be between 1-5% worldwide, constituting a serious global public health concern (Roozen et al., 2016). However, alcohol is not the only drug consumed by women during pregnancy. Cannabis is the most common illicit drug consumed by pregnant women; prevalence rates of cannabis use during pregnancy are estimated to be between 3-10% in the U.S., with some variability dependent on the legal status of cannabis across states (Coleman-Cowger et al., 2017; Ko et al., 2015; Obisesan et al., 2020; Oh et al., 2017; Samhsa, 2019; Volkow et al., 2019; Young-Wolff et al., 2019). In fact, some women purposefully use cannabis while pregnant to combat symptoms such as nausea and pain (Dickson et al., 2018), although recent reports suggest that cannabis use can actually provoke nausea and vomiting (Kim et al., 2018). These use rates and patterns are undoubtedly fueled by the perception that cannabis use is safe (Brown et al., 2017; Johnston et al., 2015), even during pregnancy (Jarlenski et al., 2017).

Despite the high rates of use, the potential consequences of prenatal cannabis exposure on the developing fetus are not well understood, as results from both clinical and preclinical studies have been mixed (Huizink, 2014; Huizink & Mulder, 2006). Results vary greatly based on developmental timing of exposure, the type of cannabinoid (phyto- or syntho-cannabinoid), the dose of cannabinoid, the type of control groups used, and the age and domain of behavior testing (Huizink, 2014; Schneider, 2009). Variability is likely to be even more enhanced given the more recent selective cultivation of cannabis constituents, which has led to a wider variety in concentrations of delta-9-tetrahydrocannabinol (THC), the primary psychoactive component. For example, the average potency of THC in cannabis products has risen from 3.4% in 1993 to 17.1% in 2019 (Chandra et al., 2019); thus, longitudinal clinical studies examining the consequences of prenatal cannabis exposure in individuals exposed decades ago may not fully inform the consequences of current potency levels consumed today (Gunn et al., 2016; Huizink, 2014; Jaddoe et al., 2012; Paul et al., 2019). Additionally, researchers are now more aware of the importance of properly assessing subtleties in neurological and behavioral alterations following prenatal cannabis exposure, even at lower doses. Clinically, the long-term consequences of cannabis products on the market today will not be known for years to come.

In addition to changes in potency and legal status, the routes of administration for cannabis use have drastically changed. The use of electronic cigarettes (e-cigarettes) for cannabis consumption has increased in popularity and more easily allows for customized combinations and potency of cannabis constituents, such as THC. Estimated rates of general e-cigarette use (vaping) among pregnant women is 5-14% (Cardenas et al., 2019), which includes cannabis vaping. This high rate is driven by the assumption that using e-cigarettes is safer than traditional smoking (Mark et al., 2015). However, consuming drugs, like cannabis, via e-cigarettes leads to higher drug and metabolite levels in the blood in both the consumer and the fetus than traditional routes (Young-Wolff et al., 2020). Thus, the potential consequences of prenatal cannabis exposure via vaping may be even more detrimental than other routes of administration. Medical professionals have requested research addressing whether prenatal e-cigarette use leads to more severe health consequences (Brandon et al., 2015; Suter et al., 2015), yet research is still in its infancy.

Moreover, polydrug consumption is also seen among pregnant women; for example, half of women who report using cannabis during their pregnancy also report consuming alcohol (Samhsa, 2015). Rates of alcohol and cannabis co-consumption among pregnant women are likely underestimated, given that self-report may be influenced by stigmatization and/or legal repercussions (Lange et al., 2014; Young-Wolff et al., 2017). To date, few studies have investigated the effects of combined alcohol and cannabinoid exposure (Abel & Subramanian, 1990; Abel et al., 1987; Basavarajappa et al., 2008; Goldschmidt et al., 2004; Hansen et al., 2008; Nagre et al., 2015; Subbanna et al., 2013). Thus, research examining possible consequences of combined alcohol and cannabis exposure during pregnancy is desperately needed.

Prenatal cannabis and alcohol exposure may influence development in many behavioral domains, including motor function. Research on the effects of prenatal cannabis on motor skill development and coordination have been mixed. On the one hand, prenatal THC exposure has been shown to decrease early developmental motility (Fried, 1976); similarly, prenatal exposure to a synthetic cannabinoid (WIN 55,212-2) impairs motor coordination later in life (Shabani et al., 2011). However, research from our lab has shown that exposure to the synthetic cannabinoid (CP-55,940) during the 3^rd^ trimester equivalent can advance early motor development, but not affect motor coordination later in life (Breit et al., 2019a). Results from clinical research are similarly conflicting. Consistent with our preclinical findings, one study found that prenatal cannabis exposure advanced motor behaviors in children 3-4 years of age (Fried & Watkinson, 1990). In contrast, others report that prenatal cannabis exposure impairs motor skills at 9 months of age, although no effects were seen at 1 year of age (Richardson et al., 1995). Additionally, motor impairments have been observed in 1-year-old children exposed to cannabis postnatally, through breastmilk (Astley & Little, 1990).

Consistent with changes in motor function, neurological alterations in motor-related areas have also been observed following prenatal THC exposure. Prenatal THC exposure in mice is associated with impaired fine motor skills and decreased cannabinoid type 1 (CB1) receptor functioning in corticospinal neurons (De Salas-Quiroga et al., 2015). Furthermore, when THC is administered during gastrulation, zebrafish showed reduced locomotor responses to sound, with altered synaptic activity at neuromuscular junctions and motor neuron morphology (Ahmed et al., 2018). Clinical research examining the effects of prenatal cannabis on motor-related brain areas has not yet been reported.

Importantly, cannabis exposure during early development may also influence overall activity levels. Increased activity levels have been observed in rodent studies following neonatal CP-55,940 exposure (Breit et al., 2019b), prenatal WIN 55,212-2 exposure (Mereu et al., 2003), and prenatal THC exposure (Borgen et al., 1973; Navarro et al., 1994; Rubio et al., 1995). However, other preclinical research has not identified any changes in general locomotor activity following prenatal THC exposure (Brake et al., 1987; Hutchings et al., 1989; Vardaris et al., 1976). Similar inconsistencies are seen in activity levels of children exposed to cannabis prenatally. While some clinical research has reported hyperactivity following prenatal cannabis exposure at 6 (Fried et al., 1992) and 10 years of age (Goldschmidt et al., 2000), other data suggest that 1-2 year old children prenatally exposed to cannabis showed no change in activity (Fried & Watkinson, 1988). Thus, it is relatively unclear how prenatal cannabis exposure alters motor-related behaviors later in life, likely due to the methodological and other differences across these studies.

In contrast to the inconsistencies seen among the cannabis literature, it has been well established that prenatal alcohol exposure impairs motor skills. Individuals with FASD consistently show impaired fine and gross motor skills (Connor et al., 2006; Driscoll et al., 1990; Kalberg et al., 2006). Importantly, preclinical studies administering alcohol during early development show similar impairments in motor development and coordination (Driscoll et al., 1990; Idrus et al., 2017; Thomas et al., 2009; Thomas et al., 2004). Moreover, developmental alcohol exposure is also often associated with attention deficit hyperactivity disorder symptom presence in clinical populations (Nash et al., 2006; O’malley & Nanson, 2002; Popova et al., 2016) and similarly leads to hyperactivity in rodents (Bond, 1981; Breit et al., 2019b; Ryan et al., 2008; Thomas et al., 2007).

Research examining the behavioral effects of combined alcohol and cannabinoid exposure during the gestational period is limited, despite the high prevalence of co-consumption of alcohol and cannabis during pregnancy. The few studies that have explored alcohol and cannabinoid exposure during early development have primarily focused on the effects of either drug individually or combined effects on physical, physiological, or neural pathologies (Abel et al., 1987; Fish et al., 2019; Hansen et al., 2008). Our lab recently examined the effects of combined exposure to alcohol and cannabinoids during the third trimester equivalent. We illustrated that combined neonatal exposure to CP-55,940 and alcohol impaired motor coordination among adolescent females (Breit et al., 2019a). Similarly, although neonatal exposure to alcohol and CP-55,940 separately increased locomotor activity among adolescent offspring, the combination produced even more severe hyperactivity, in part by reducing habituation (Breit et al., 2019b). These results suggest that combined exposure to alcohol and THC during early development may have more severe effects on motor development, coordination, and activity than either drug alone. Moreover, combined exposure where THC is administered via e-cigarette may be even more detrimental (Young-Wolff et al., 2020), as combined exposure may alter drug metabolism, increasing individual drug and metabolite levels in the blood of the user in both clinical (Downey et al., 2013; Hartman et al., 2015) and preclinical research (Breit et al., 2020).

To address this question, the current study examined whether prenatal exposure to vaporized alcohol, THC via e-cigarette, or the combination would alter early sensorimotor development, later motor coordination and activity levels among offspring. We used a novel vapor inhalation model using commercially available e-cigarette exposure. Importantly, our polydrug vapor inhalation model intoxicates dams with alcohol and/or THC while avoiding confounding nutritional factors, as previously established (Breit et al., 2020). Prenatal vaporized drug exposure occurred once daily during a period equivalent to the human first and second trimesters. Following birth, offspring were tested on a grip strength/ hindlimb coordination task early in development, a parallel bar motor coordination task during adolescence, and adolescent open field activity to examine potential effects of prenatal exposure on motor behaviors across development.

## 2.1 Materials and Methods

### 2.1.1 Maternal Vapor Exposure

#### 2.1.1.1 Dams

All procedures and behavioral tests included in this study were approved by the San Diego State University Institutional Animal Care and Use Committee and are in accordance with the National Institute of Health’s *Guide for the Care and Use of Laboratory Animals*.

Naïve female Sprague-Dawley rats were obtained from Charles River Laboratories (Hollister, CA) and allowed to acclimate for 2 weeks before any procedures began. Prior to breeding, intravenous catheters were surgically implanted to measure blood alcohol and THC levels during pregnancy (details described in Breit et al. 2020). Following intravenous catheter placement, each dam was housed with a stud for up to 5 consecutive days. The presence of a seminal plug was deemed as gestational day (GD) 0; pregnant dams were then singly housed and monitored daily until the day of birth (typically GD 22, also known as postnatal day [PD] 0). On GD 0, dams were also randomly assigned to 1 of 4 prenatal vapor inhalation exposure groups: the combination of ethanol (EtOH) and THC (EtOH+THC), EtOH alone (EtOH+Vehicle), THC alone (Air+THC), or controls (Air+Vehicle).

#### 2.1.1.2 Prenatal Vapor Inhalation

The vapor inhalation equipment (La Jolla Alcohol Research Inc, CA) used both heated flask vaporization of EtOH and e-cigarette atomization for THC and Vehicle exposure. From GD 5-20, dams were exposed once daily to ethanol (EtOH; 95%, Sigma Aldrich) or Air via vapor inhalation (10 L/min) for 3 hours in a constant stream. Following EtOH or Air exposure, dams were then exposed to THC (100 mg/mL; NIDA Drug Supply Program) or the Vehicle (propylene glycol, Sigma Aldrich) via an e-cigarette tank (SMOK V8 X-Baby Q2) for 30 minutes (2 L/min). E-cigarette exposure to THC or the Vehicle was delivered in seven 6-sec puffs that were 5-minutes apart (30 min total), followed by 10 min of air exposure to clear out any residual vapors in the chambers.

Following birth, on PD 2, litters were culled to 4 females and 4 males (when possible). To avoid potential litter effects, only one sex pair (1 female, 1 male) per litter was tested on the early sensorimotor development and motor coordination tasks; a separate sex pair was examined in open-field activity chambers. Remaining offspring were used for other behavioral tasks, which will be reported elsewhere. On PD 7, offspring were tattooed with non-toxic veterinary ink for identification purposes. Litters were weaned from the dam on PD 21 and separated by sex on PD 28. All motor testing took place between PD 12-34 (Figure 1). Subjects had *ad libitum* access to food and water, were weighed daily, and were acclimated to the testing room 30 min prior to each task.

**Figure 1.**
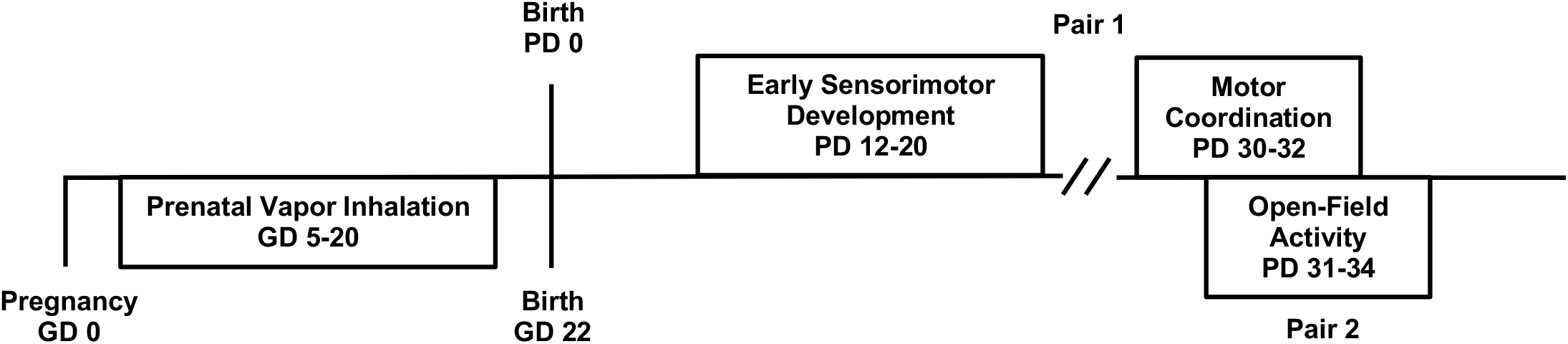
Timeline of the study design.

### 2.1.2 Early Sensorimotor Development

Offspring completed a grip strength and hindlimb coordination test daily from PD 12-20 to examine early sensorimotor development. Pups were placed with their forepaws on a wire (2-mm diameter) suspended above a cage of bedding. Each subject was given 2 consecutive trials (30 sec each) to either hold onto the wire for the full 30 sec or place their hindlimb on the wire; if either were achieved, the trial was recorded as successful. The first day of any success, and the number of successes were recorded live by two investigators; all testing was also recorded for additional analysis.

### 2.1.3. Adolescent Motor Coordination

The same offspring examined on early sensorimotor development were later tested for motor coordination and balance using a parallel bar task daily from PD 30-32. The apparatus included two parallel steel bars (0.5 cm diameter, 91 cm length) bolted between two platforms (15.5 cm × 17.8 cm) above a cage of bedding. The platforms have grooved slots (0.5 cm apart), allowing the width between the bars to be altered at 0.5-cm increments. The parallel bar testing was conducted in a room illuminated with a red light to promote locomotor activity.

Offspring were placed on one of the two platforms for 30 sec to acclimate to the apparatus. Subjects were then placed in the middle of the parallel bars, with one forelimb and one hindlimb on each parallel bar. Once subjects were able to successfully traverse across the bars with four consecutive hindlimb steps, the trial was deemed successful. If subjects’ paws slipped, they fell, or they swung below the bars, the trial was considered a failure. With each successful trial, the width of the parallel bars was increased by 0.5 cm. Subjects were given up to 15 trials per day for 3 consecutive days. Testing ceased for the day once the maximum number of trials were reached (15), or if the subject failed on 5 consecutive trials. The trials to the first success, number of successful trials, maximum width achieved, and the success ratio (successful trials divided by total trials) were recorded each day.

### 2.1.4. Adolescent Activity Levels

Activity levels were examined using an open-field activity chamber paradigm from PD 31-34; offspring tested in this domain were separate sex pairs from those tested on early sensorimotor development and motor coordination. Activity levels were assessed during the offspring’s dark cycles (beginning at 18:00). Each subject was placed in an individual automated open-field activity chamber with infrared beams (16 × 16 × 14 in; Hamilton-Kinder) for 60 min; chambers were equipped with white noise and fans for ventilation. The number of infrared beam breaks, distance traveled (in), number of rears, entries into the center, and time spent in the center were recorded in 5-minute time bins for each day of testing.

### 2.1.5. Statistical Analyses

Offspring motor data were analyzed using the Statistical Package for Social Sciences (IBM; version 27). All data were initially tested for normality (Shapiro-Wilk test). Data that were normally distributed were analyzed using a 2 (EtOH, Air) × 2 (THC, Vehicle) × 2 (Female, Male) design, with repeated measures (Day, Bin) when necessary. Binomial data, data with a limited range of outcomes, or data that were not normally distributed were analyzed non-parametrically using Mann-Whitney, Kruskal-Wallis, or Fishers-exact analyses, where appropriate. Means (M) and standard error of the means (SEM) are reported when applicable. Significance levels for all paradigms were set at *p <* 0.05.

## 3.1 RESULTS

### 3.1.1. Body Weights

#### 3.1.1.1. Motor Development

A total of 88 subjects completed both the early sensorimotor and the motor coordination tasks (EtOH+THC: 9 females, 8 males; EtOH+Vehicle: 13 females, 10 males; Air+THC: 13 females, 11 males; Air+Vehicle: 12 females, 12 males). Body weights for all tasks were analyzed with a EtOH × THC × Sex ANOVA with Day as a repeated measure. As previously published, neither litter size nor initial birth weights differed among any of the prenatal exposure groups (Breit et al., 2020).

During sensorimotor testing, subjects prenatally exposed to THC weighed less than those not exposed to THC, producing both a significant main effect of THC (F[1,80] = 19.520, *p =* 0.00003) and an interaction of THC × Day (F[8,640] = 7.854, *p =* 4.2284^E-10^). Although offspring in all groups gained significant weight during this developmental period, THC-exposed offspring weighed less than Vehicle-exposed offspring on each Day (all *p*’s < 0.001 for PD 12-20), and the THC-related reductions were more pronounced as days progressed (Figure 2A). Thus, prenatal exposure to THC via e-cigarettes reduced offspring body weights during this early period. In contrast, prenatal EtOH exposure did not alter body weights during this developmental period, nor were there any significant main or interactive effects of Sex.

**Figure 2.**
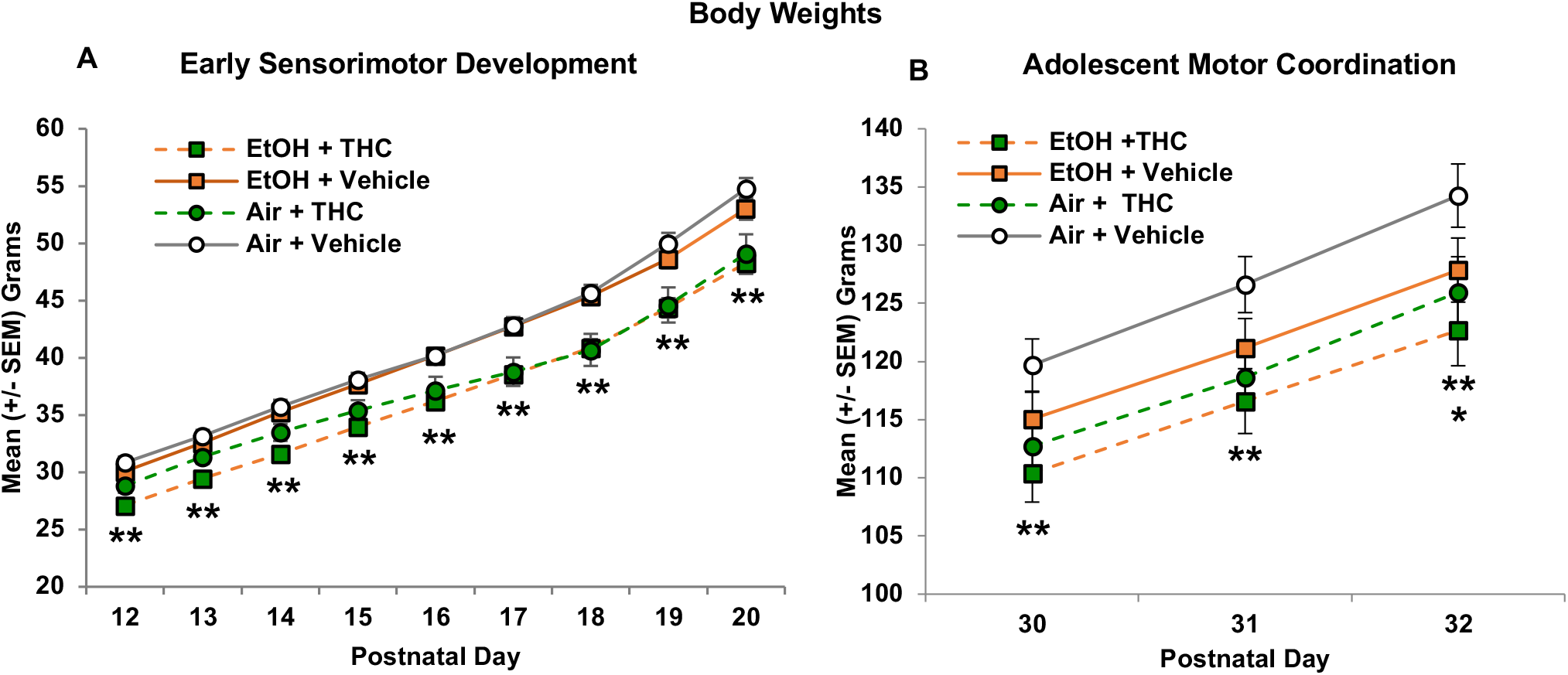
Offspring exposed to THC prenatally weighed significantly less throughout the early sensorimotor development paradigm (postnatal days 12-20; A) and the parallel bar paradigm (PD 30-32; B), alone or in combination with alcohol. Separately, there was an overall reduction in body weight of alcohol-exposed subjects on PD 32 (B), an effect driven by the combined exposure group. ** = THC < Vehicle, *p*’s < 0.05. * = EtOH < Air, *p =* 0.05.

Prenatal THC exposure continued to reduce body weight throughout parallel bar testing (PD 32). Offspring exposed to THC prenatally weighed less than their Vehicle-exposed counterparts overall (F[1,79] = 6.338, *p =* 0.0138, alone or in combination with EtOH (Figure 2B). In addition, a Day*EtOH (F[2,158] = 3.428, *p =* 0.0349) was observed. Offspring exposed to EtOH prenatally weighed less than their Air-exposed counterparts on the last day of testing (PD 32; F[1,79] = 3.86, *p =* 0.0531), although this effect was driven by the combined exposure group (F[3,83] = 2.637, *p =* 0.0551; SNK *p <* 0.05; Figure 2B), as body weights of subjects exposed to EtOH only did not significantly differ from that of controls. Overall, females weighed less than males (F1,79] = 30.291, *p =* 4.4618E^−7^) and gained weight more slowly than males over these days, leading to a Day*Sex interaction (F[2,158] = 23.539, *p =* 1.1277E^−9^); however, Sex did not interact with EtOH or THC.

#### 3.1.1.2. Adolescent Activity Levels

Separate littermates (88 subjects) were examined for open field activity levels (EtOH+THC: 9 females, 9 males; EtOH+Vehicle: 12 females, 11 males; Air+THC: 13 females, 12 males; Air+Vehicle: 12 females, 12 males). Across Days, females weighed less than males, producing a main effect of Sex (F[1,82] = 47.236, *p =* 1.1308E^−9^), but Sex did not interact with prenatal exposure EtOH or THC. Overall, offspring exposed to THC only weighed significantly less than all other groups, producing a significant interaction of EtOH*THC (F[1,82] = 6.907, *p =* 0.0103), Day*THC (F[3,246] = 3.558, *p =* 0.0150), as well as a main effect of THC (F[1,82] = 10.871, *p =* 0.0014). Subjects exposed to only THC weighed less than all other Groups on all days (PD 31: F[3,82] = 6.331, *p =* 0.0006; PD 32: F[3,82] = 7.424, *p =* 0.0002; PD 33: F[3,82] = 7.381, *p =* 0.0002; PD 34: F[3,82] = 6.436, *p =* 0.0006; all SNK *p*’s < 0.05; Figure 3).

**Figure 3.**
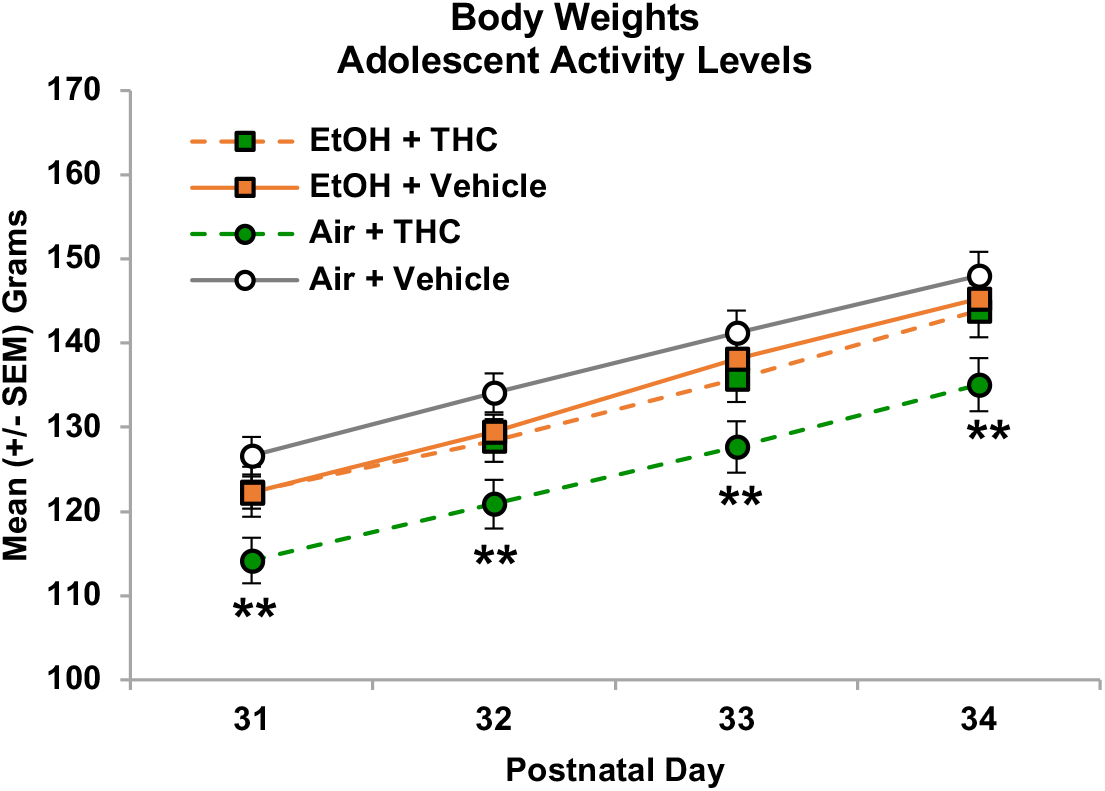
Offspring exposed to THC alone weighed less than all other prenatal exposure groups during the open-field activity testing. ** = Air+THC < all other groups, *p*’s < 0.05.

In addition, separate 2-way interactions of Day*EtOH (F[3,246] = 3.387, *p =* 0.0188) and Day*Sex (F[3,246] = 44.812, *p =* 3.8574E^−23^) were seen in the body weight data from PD 31-34. However, there were no statistically significant effects of EtOH on any day and follow-up analyses simply confirmed that females consistently weighed less than males on each day (PD 31: F[1,82] = 32.195, *p =* 2.0376E^−7^; PD 32: F[1,82] = 42.199, *p =* 5.9711E^−9^; PD 33: F[1,82] = 49.937, *p =* 4.7631E^−8^; PD 34: F[1,82] = 61.984, *p =* 1.2463E^−11^).

### 3.1.2. Early Sensorimotor Development

Shapiro-Wilk normality tests indicated that the behavioral data for this task were not normally distributed (all *p*’s < 0.05) due to the limited numerical range of outcomes. Thus, non-parametric analyses were used for all variables, as appropriate. Separate Mann-Whitney U tests comparing main effects of EtOH and THC were initially conducted. In addition, since non-parametric analyses do not allow for examination of interactions, data were also analyzed across the 4 exposure groups using Kruskal-Wallis H tests. Lastly, for binomial data (i.e. able to perform or not), Chi-Square comparisons were used. Females and males did not significantly differ on any behavioral outcome measures, so all data were collapsed by Sex.

Exposure to THC via e-cigarettes during gestation significantly delayed the age of the first successful trial on the sensorimotor task (U = 648.00, *p =* 0.0066; H = 9.850, *p =* 0.0020; Figure 4A). In contrast, prenatal EtOH exposure had no significant effect on the first day of a successful trial (U = 870.00, *p =* 0.4282). Chi-square comparisons on each Day during the early sensorimotor task indicated that prenatal THC exposure delayed successful performance on this task, both alone and in combination with EtOH (Figure 4B). On PD 12, offspring exposed to the combination of EtOH+THC prenatally performed worse than those exposed to EtOH only (EtOH+Vehicle: *X*^2^ = 3.913, *p =* 0.0484) and controls (Air+Vehicle: *X*^2^ = 8.129, *p =* 0.0050), whereas THC-exposed subjects (Air+THC) were impaired compared to control offspring (*X*^2^ = 4.090, *p =* 0.0041). Subjects exposed prenatally to the combination of EtOH+THC continued to perform worse than those exposed to EtOH only (*X*^2^ = 4.365, *p =* 0.0039) and controls (*X*^2^ = 4.784, *p =* 0.0031) on PD 13, but by PD 15, no differences were seen among Groups, suggesting a catch-up in sensorimotor development.

**Figure 4.**
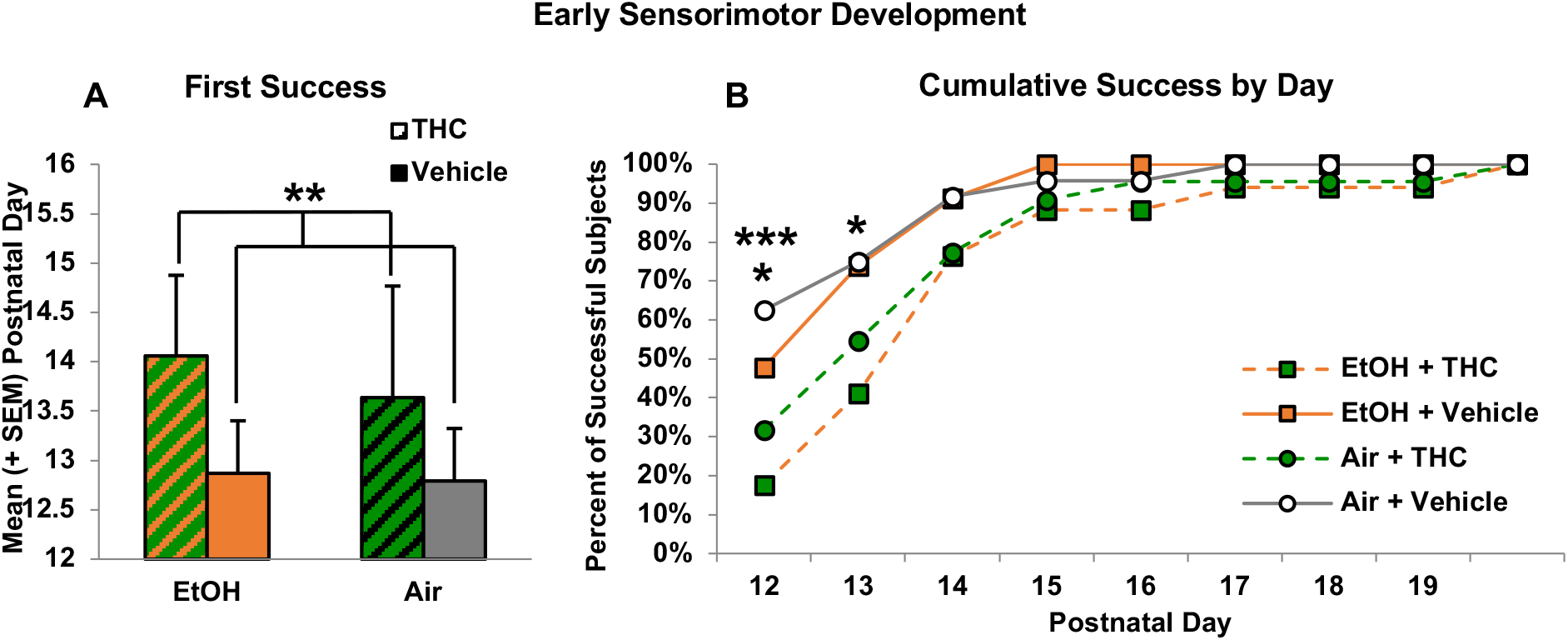
Prenatal exposure to THC via e-cigarettes delayed early sensorimotor development. Although subjects exposed to prenatal THC showed developmental delays (A), their motor development eventually caught up with that of controls (B). ** = THC > Vehicle, *p <* 0.05. * = EtOH+THC < EtOH+Vehicle and Air+Vehicle, *p*’s < 0.05. *** = Air+THC < Air+Vehicle, *p <* 0.05.

### 3.1.3. Adolescent Motor Coordination

Prenatal THC exposure also significantly impaired performance on all measures of the parallel bar motor coordination task, indicating that in addition to early delays in motor development, prenatal THC leads to long-lasting motor dysfunction. Offspring exposed to THC prenatally required more trials to reach their first success, producing a main effect of THC (F[1,68] = 7.237, *p =* 0.0090; Figure 5A). Similar effects of prenatal THC exposure were seen on the success ratio of the first day of testing, as subjects exposed to prenatal THC were significantly less successful compared to groups not exposed to THC (F[1,68] = 10.379, *p =* 0.0020; Figure 5B), consistent with the percent of subjects successful on day 1 (EtOH+THC: 41%, EtOH+Vehicle: 63%, Air+THC: 43%, Air+Vehicle: 65%). Interestingly, prenatal alcohol exposure did not affect these outcomes, nor did it modify the effects of prenatal THC exposure.

**Figure 5.**
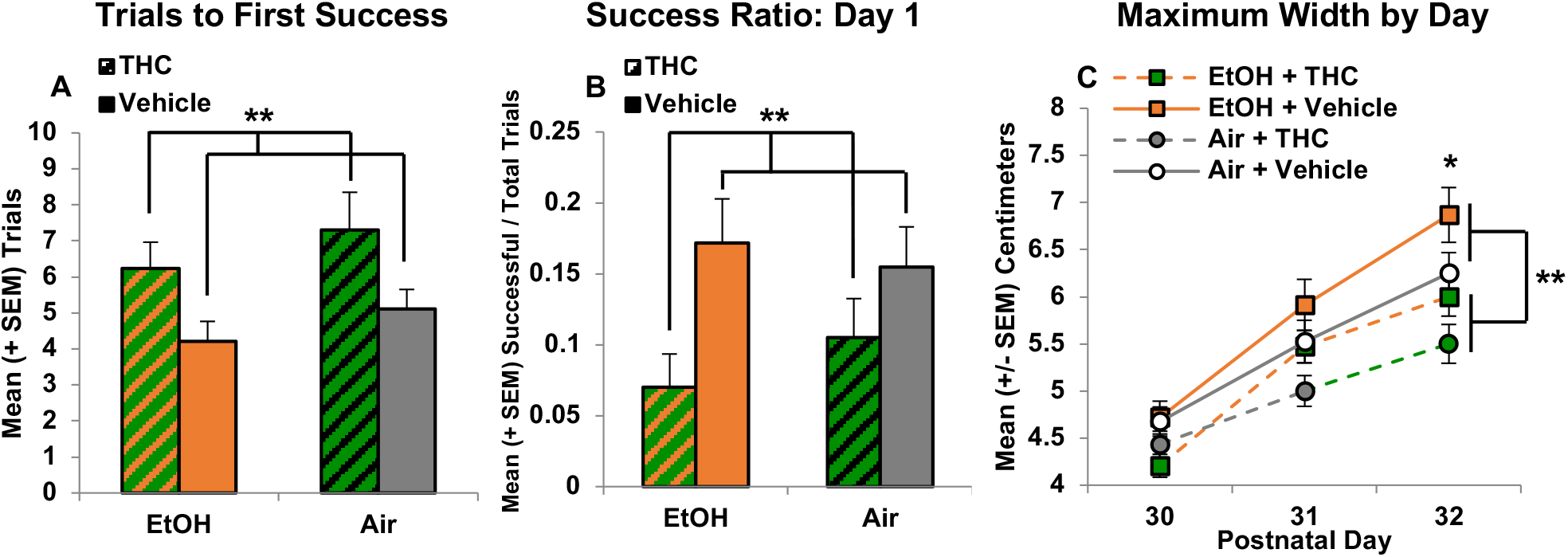
Prenatal THC exposure impaired motor coordination in adolescent subjects, increasing the number of trials before the first success (A), decreasing success ratios (B), and decreasing the maximum width achieved (C). ** = THC significantly different from Vehicle *p*’s < 0.05. * = EtOH > EtOH+THC and Air+THC, *p <* 0.001.

Finally, prenatal THC exposure also reduced the maximum width successfully traversed (F[1,68] = 12.165, *p =* 0.0013; Figure 5C). As expected, performance of subjects improved over days, producing a main effect of day (F[2,136] = 173.241, *p =* 4.0176E^−38^). However, there was also a significant interaction of Day*EtOH (F[2,136] = 7.896, *p =* 0.0006), as EtOH-exposed offspring collapsed across groups were able to traverse greater widths compared to their Air-exposed counterparts on the last day of testing (PD 32; F[1,68] = 5.135, *p =* 0.0266; Figure 5C); there were no significant differences between the EtOH only and controls. Male and female offspring did not differ in any measure of motor coordination, nor did Sex interact with prenatal EtOH or THC exposure.

### 3.1.4. Open Field Activity Levels

#### 3.1.4.1. Locomotor Activity

In contrast to motor development, activity levels were most affected by the combination of EtOH and THC in a sex-dependent manner. The total distance traveled in the open field declined across Days (F[3,246] = 20.509, *p =* 6.787E^−12^) and Time (F[11,902] = 345.702, *p =* 0.0E^0^) within each session. Although there were no main effects of EtOH, THC, Sex, there were significant interactions of Day*EtOH*THC (F[3,246] = 5.977, *p =* 0.0006), Bin*THC (F[11,902] = 1.944, *p =* 0.0310), and a trending 5-way interaction of Day*Bin*Sex*EtOH*THC (F[33,2706] = 1.400, *p =* 0.0647). Very different patterns were evident between female and male offspring; thus, follow-up analyses were conducted separately for each Sex.

Among female offspring, separate interactions of Day*Bin*EtOH*THC (F[33,1386] = 1.525, *p =* 0.0293) and Bin*EtOH*THC (F[11,462] = 2.013, *p =* 0.0257) were observed. Follow-up analyses suggest that these interactions were due to subtle increases in activity levels among females exposed to either prenatal EtOH or prenatal THC exposure compared to controls (Air+Vehicle) during the middle of the testing sessions; Figure 6B).

**Figure 6.**
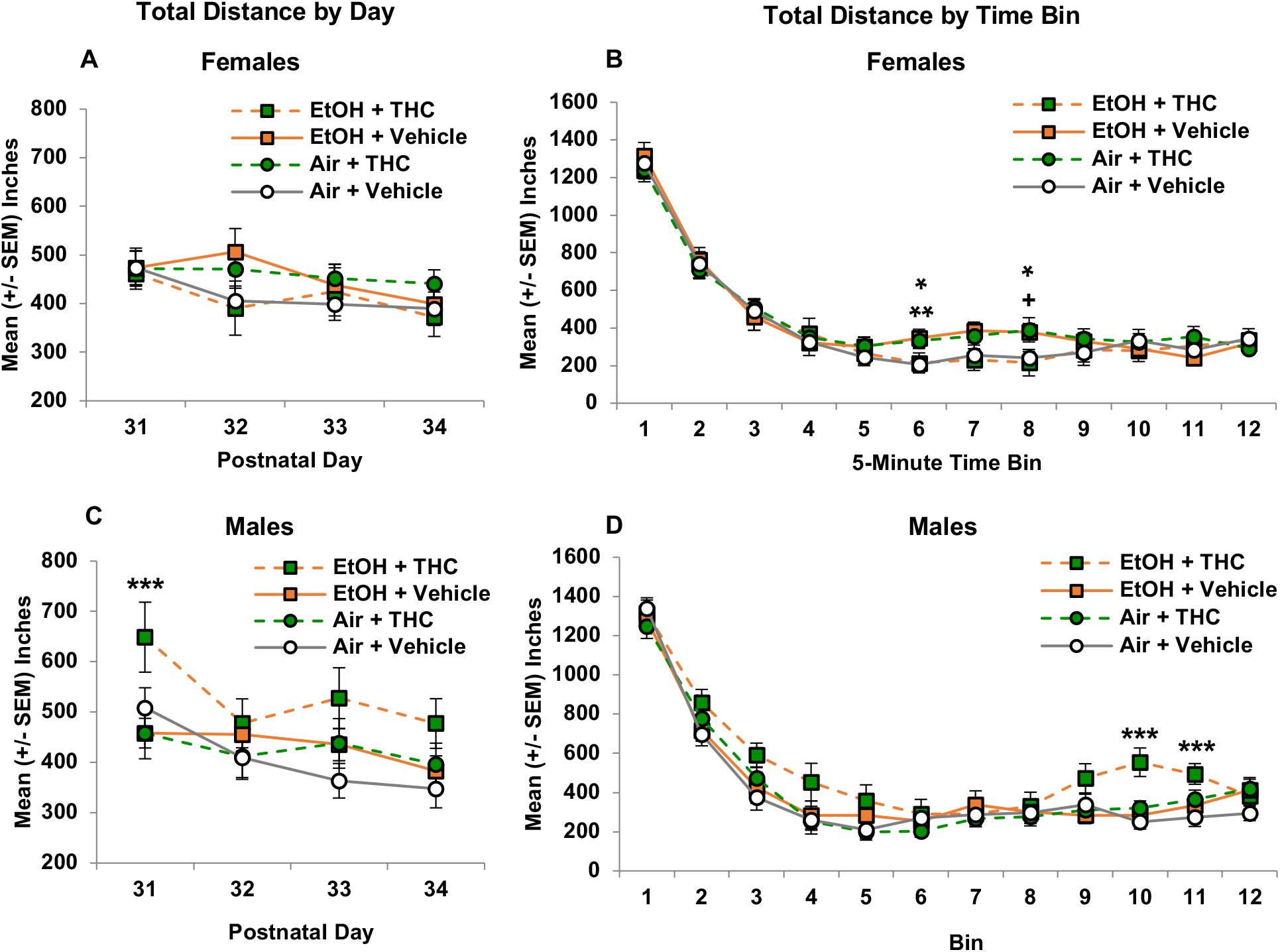
Among female offspring, prenatal EtOH or THC exposure separately increased activity levels during the middle of the sessions compared to controls (B). In contrast, combined exposure to prenatal EtOH and THC significantly increased activity levels among male offspring overall on Day 1 (C), as well as towards the end of the testing sessions (D). * = EtOH+Vehicle > Air+Vehicle, *p*’s < 0.05. ** = Air+THC > Air+Vehicle, *p <* 0.05. + = Air+THC > Air+Vehicle, *p =* 0.06. *** = EtOH+THC > all other groups, *p*’s < 0.05.

In contrast, among male offspring, the combination of prenatal EtOH and THC significantly increased locomotor activity, producing significant interactions of Day*EtOH*THC (F[3,120] = 6.017, *p =* 0.0007) and Bin*THC (F[11,440] = 2.314, *p =* 0.0091). Follow-up analyses indicated that male offspring exposed to the combination of EtOH+THC traveled significantly further on the first day of testing (PD 31; F[3,40] = 3.278, *p =* 0.0307; all SNK *p*’s < 0.05; Figure 6C). Within sessions, male offspring prenatally exposed to EtOH+THC also traveled farther than the other Groups toward the end of the sessions (Bin 10: F[3,40] = 7.879, *p =* 0.0003; Bin 11: F[3,40] = 3.102, *p =* 0.0373; all SNK *p*’s < 0.05; Figure 6D). Similar group patterns were seen in distance traveled in the periphery of open field, fine movement, and overall number of beam breaks (data not shown).

#### 3.1.4.2. Center Distance

Effects of prenatal drug exposures were particularly striking when examining locomotor activity within the center of the chamber (inner 8 × 8 inches). As expected, the center distance traveled declined across Days (F[3,246] = 20.885, *p =* 4.3539E^−12^) and Time within each session (F[11,902] = 244.424, *p =* 1.2231E^−261^). There were also significant interactions of Sex*EtOH*THC (F[1,82] = 5.284, *p =* 0.0241), Day*EtOH*THC (F[3,246] = 4.782, *p =* 0.0029), and Bin*Sex (F[11,902] = 1.830, *p =* 0.0453).

Among female offspring, no further interactions or main effects of any variable reached significance (Figures 7A and 7B). In contrast, among male offspring, there was an overall interaction of EtOH*THC (F[1,40] = 4.180, *p =* 0.0475), as males exposed to the combination of EtOH and THC prenatally traveled significantly further in the center of the chamber compared to all other groups. There was also an interaction of Day*EtOH*THC (F[3,120] = 4.369, *p =* 0.0059); the combination of EtOH and THC exposure increased locomotor activity in the chamber center on the first (F[1,40] = 5.377, *p =* 0.0033; all SNK *p*’s < 0.05) and third (F[3,40] = 3.111, *p =* 0.0369; all SNK *p*’s < 0.05) days of testing (Figure 7C), failing to reach significance on the fourth day (F[1,40] = 2.489, *p =* 0.0742). Within sessions, males exposed to the combination traveled more compared to all other groups early in the sessions (Bins 3-5; all *p*’s < 0.05) and toward the end of the sessions (Bins 9-11; all *p*’s < 0.05; Figure 7D). Similar effects were seen with the number of center entries (data not shown).

**Figure 7.**
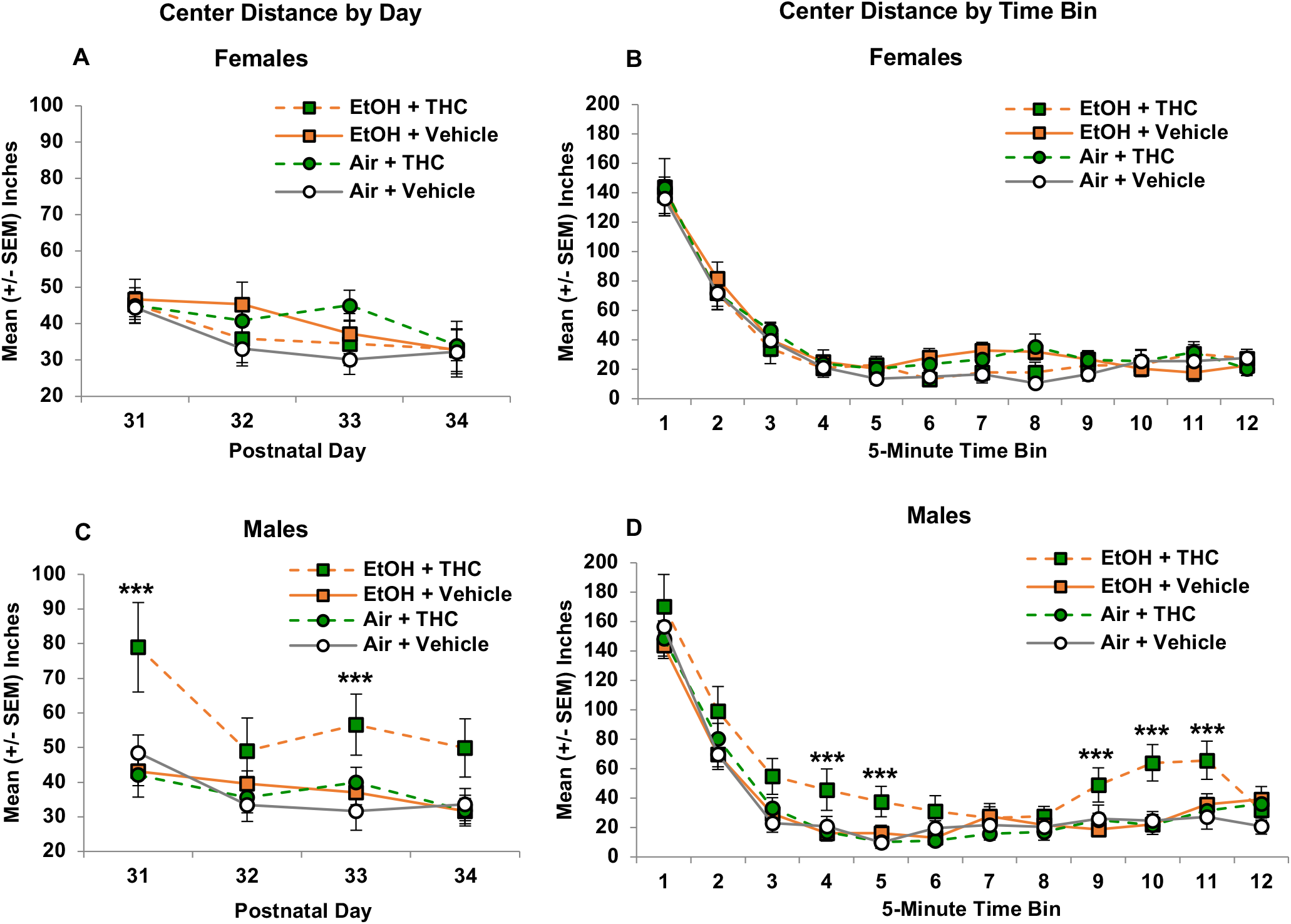
Prenatal exposure did not significantly change the distance traveled in the center of the chamber among females on any day of activity testing (A) or across testing sessions (B). However, male offspring exposed to the combination of alcohol and THC prenatally traveled greater distances on Days 1 and 3 of testing (C) and throughout the testing session durations (D), indicating impaired habituation. *** = EtOH+THC > all other groups, *p <* 0.05.

#### 3.1.4.3. Center Time

As expected, males exposed to the combination of EtOH and THC also spent more time in the center of the chamber. Time spent (sec) in the center of the chamber decreased across Days (F[3,246] = 2.875, *p =* 0.0368) and Time Bins (F[11,902] = 52.644, *p =* 1.8843E^−89^) within each session. In addition, interactions of Sex*EtOH*THC (F[1,82] = 8.401, *p =* 0.0048), Sex*EtOH (F[1,82] = 3.700, *p =* 0.0588), Day*Bin*THC (F[33,2706] = 1.558, *p =* 0.0225) and Bin*Sex (F[11,902] = 2.842, *p =* 0.0011) were evident.

Among female offspring, there were no main effects of prenatal EtOH or THC exposure (Figures 8A and 8B). In contrast, male offspring exposed to the combination of EtOH and THC spent more time in the center of the chamber overall, producing an interaction of EtOH*THC (F[1,40] = 7.351, *p =* 0.0098). There were no significant alterations with Day and Bin among males, likely due to the high amount of variability among male offspring prenatally exposed to combined EtOH+THC; however, increased center time in this group was most robust on Day 1 (F[3,40] = 5.542, *p =* 0.0028; all SNK *p*’s < 0.05; Figure 8C) and toward the end of the testing sessions (Bins 5, 9-11: all SNK *p*’s < 0.05; Figure 8D).

**Figure 8.**
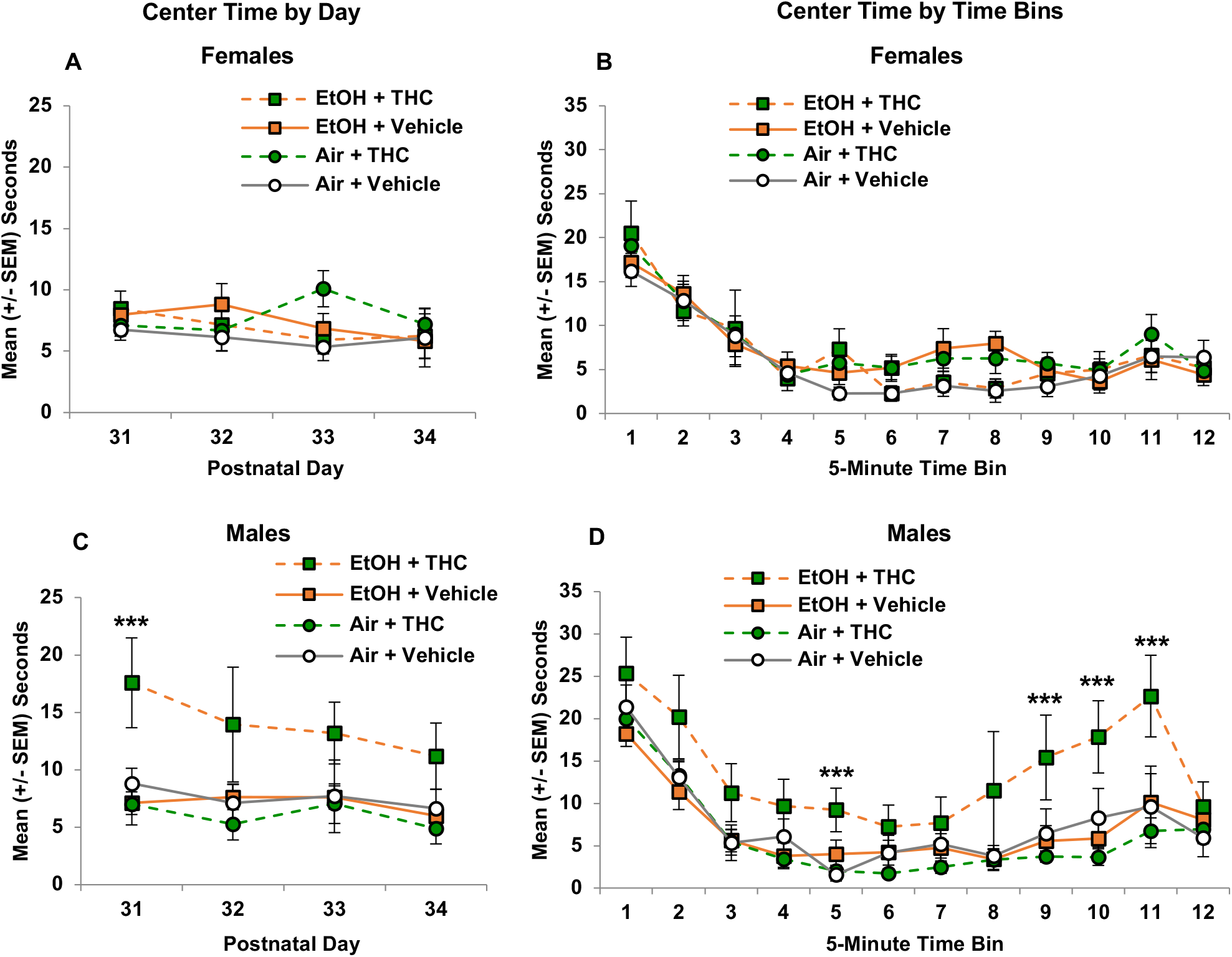
Prenatal exposure to alcohol and/or THC did not significantly alter the time spent in the center of the chamber among female offspring (A and B). However, combined exposure to alcohol and THC generally increased the center time of male offspring in the open field on Day 1 (C) and at the end of the testing sessions (D). *** = EtOH+THC > all other groups; *p*’s < 0.05.

#### 3.1.4.4 Rearing

In contrast to locomotor activity, no main or interactive effects of Sex were observed on rearing, an exploratory behavior. The number of times subjects reared (standing on hindlimbs) within the chamber declined across Days (F[3,246] = 7.941, *p =* 0.00005) and Time Bins (F[11,902] = 243.006, *p* = 8.6475E^−261^) within each session. Although interactions of Day*EtOH*THC (F[3,246] = 4.629, *p =* 0.0036), there were no significant main or interactive effects of prenatal drug exposure when analyzing data on individual days (Figure 9A). Within sessions, offspring exposed to EtOH prenatally reared less at the very beginning of the sessions (Bin 1) than Air-exposed groups (F[1,86] = 4.288, *p =* 0.0404; Figure 9B), contributing to a Bin*EtOH*THC (F[11,902] = 2.308, *p =* 0.0086) interaction; however, no further significant effects of prenatal exposure were observed throughout the duration of the sessions.

**Figure 9.**
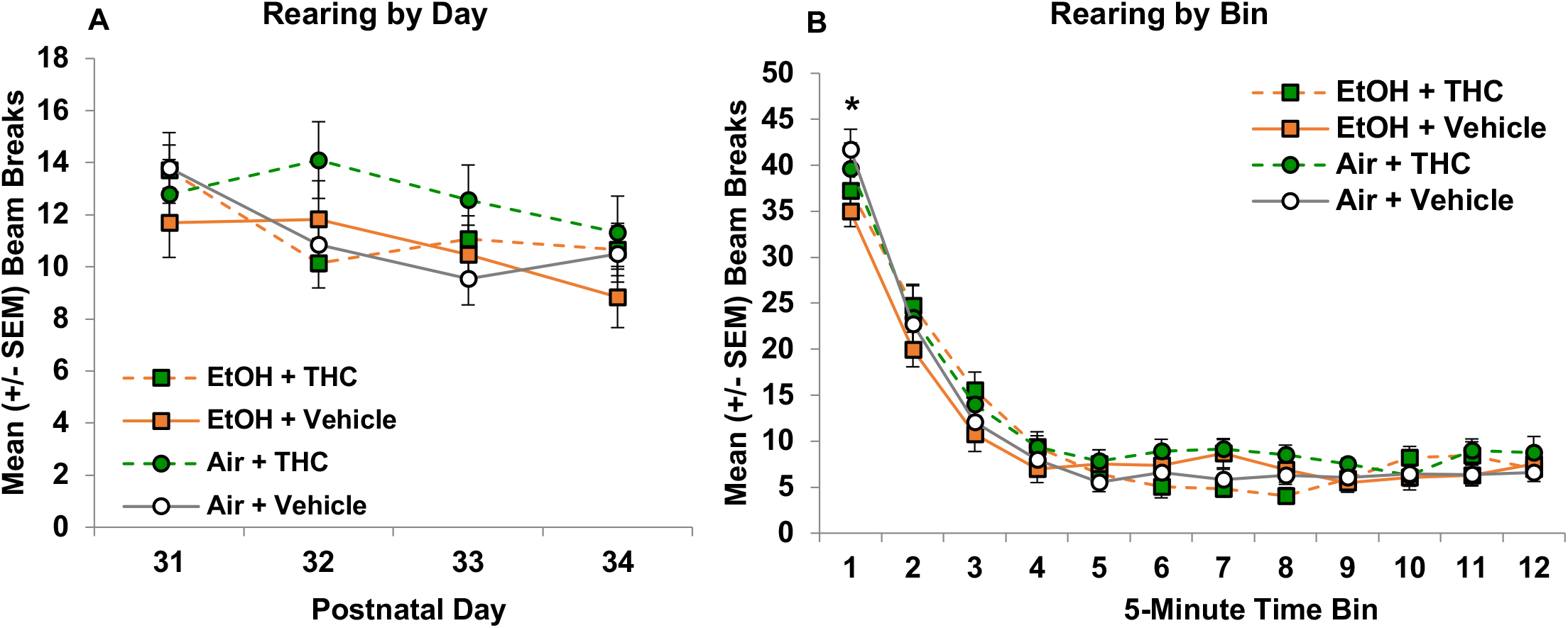
Prenatal exposure did not significantly change the frequency of rearing in the open field chamber among either sex (A). All offspring exposed to alcohol prenatally reared less than those exposed to air at the beginning of the session (B). * = EtOH < Air, *p <* 0.05.

## 4.1 DISCUSSION

This study is the first to illustrate that prenatal alcohol and THC exposure via e-cigarettes leads to both separate and interactive alterations in motor function among offspring. Prenatal THC exposure via e-cigarettes reduced body weights during both early (PD 12-20) and adolescent (PD 30-34) periods among offspring of both sexes. In addition, prenatal THC exposure also delayed early sensorimotor development and continued to impair motor coordination during adolescence, alone or in combination with alcohol. In contrast, prenatal alcohol exposure produced subtle and transient reductions in exploratory behaviors in the open-field activity chambers and both prenatal alcohol and THC alone induced modest increases in locomotor activity among females. However, only the combination of prenatal alcohol and THC exposure increased locomotor activity levels among male adolescent offspring.

Importantly, we have established that this novel, co-exposure model of vaporized alcohol and THC e-cigarettes induces physiologically relevant drug levels in the blood of pregnant rats (alcohol: 200 mg/dl; THC: 20 ng/mL), while avoiding maternal nutritional confounds or altering litter outcomes such as gestational length, litter size, litter sex ratios, or offspring birth weights (Breit et al., 2020). The doses used mimic a moderate binge dose of alcohol (twice the legal limit of 80 mg/dl) and a low-moderate dose of THC (Andrenyak et al., 2017). Given the lasting effects on offspring behavior with these THC doses, the results are particularly concerning. Interestingly, combined exposure to alcohol and THC increased maternal blood levels of each drug more than either drug alone, suggesting pharmacokinetic interactions of the combination of exposures among pregnant dams, an effect similar to clinical findings following co-consumption (Downey et al., 2013; Hartman et al., 2015). Despite this pharmacokinetic interaction, the only behavioral interactive effects in the present study were seen in open field activity among males.

Using this co-exposure vapor inhalation paradigm, physical consequences of exposure on offspring are limited. Although prenatal alcohol exposure delayed offspring eye opening (Breit et al., 2020), the only other consistent effect has been reduced body growth among offspring exposed to THC prenatally via e-cigarettes. We previously reported that prenatal THC exposure decreased body weights on PD 30 among male offspring (Breit et al., 2020); this effect was similarly seen among females, though results were not significant. The two sets of sex pairs used in the currently study are from the same litters as previously published (Breit et al., 2020). Among the sex pair tested on the motor coordination tasks, reduced body weights were observed from PD 12 to PD 32 among both male and female offspring exposed to THC prenatally, alone or in combination with alcohol. In the sex pair tested in the open field, similar body weight reductions were seen from PD 31-34 in both males and females, although only reached statistical significance among offspring exposed to prenatal THC alone. Although slight alterations in body weight reductions were seen among different sex pairs, it is important to reiterate that all offspring stemmed from the same litters, and some degree of within-litter variability is expected and relevant in teratology research (Holson & Pearce, 1992). Overall, prenatal THC exposure via e-cigarettes reduced offspring body weights during the early and adolescent periods, which is similar to findings of clinical data suggesting that prenatal cannabis exposure may impair body growth (Abel, 1980; Huizink, 2014).

In contrast, prenatal alcohol exposure via vapor inhalation did not alter body weight growth. Although prenatal alcohol exposure is associated with smaller birth weights and body growth in clinical data (Caputo et al., 2016; Carter et al., 2007; Spohr et al., 1993), alcohol-related reductions in offspring exposed prenatally are less consistent in preclinical data (Dursun et al., 2006; Hannigan et al., 1993; Helfer et al., 2009; Thomas et al., 2009; Thomas et al., 2010) and are often dose-dependent (Abel & Dintcheff, 1978). In the current study, average peak blood alcohol concentrations (BACs) among those who received alcohol alone were 159.21 ± 15.02mg/dL, whereas those in the combined group reached 205.40 ± 18.03 mg/dL (Breit et al., 2020). Previous studies who have shown reduced body weights and hyperactivity among offspring used binge-like alcohol exposure with BACs exceeding 200-300 mg/dL (Dursun et al., 2006; Thomas et al., 2009; Thomas et al., 2010).

However, impaired motor coordination following prenatal alcohol exposure in rodent models has been inconsistently seen even among research achieving these higher BAC levels (Driscoll et al., 1990; Dursun et al., 2006; Idrus et al., 2017; Thomas et al., 2009; Thomas et al., 2010). While clinical data have found prenatal alcohol-related motor impairments consistently (Connor et al., 2006; Driscoll et al., 1990; Kalberg et al., 2006), particularly among greater exposure levels (Jacobson et al., 1998), preclinical data using prenatal models have been less constant. There is some evidence that early postnatal exposure to alcohol among rodents may lead to more consistent motor impairments (Breit et al., 2019a; Thomas et al., 2004), particularly with higher doses (Kelly et al., 1987). Thus, although we expected alcohol-related motor impairments in both the early sensorimotor and later motor coordination tasks, our results are not inconsistent with other studies. In addition to the timing of exposure, it is also possible that differences in administrative route could also contribute to differences.

Although the effects of prenatal cannabinoid exposure on motor skill development have been similarly inconsistent in the literature (Astley & Little, 1990; Borgen et al., 1973; Brake et al., 1987; Breit et al., 2019a, 2019b; Fried, 1976; Fried & Watkinson, 1988; Fried & Watkinson, 1990; Fried et al., 1992; Goldschmidt et al., 2000; Hutchings et al., 1989; Mereu et al., 2003; Navarro et al., 1994; Richardson et al., 1995; Rubio et al., 1995; Shabani et al., 2011; Vardaris et al., 1976), we observed clear delays in early sensorimotor development among offspring following prenatal THC exposure via e-cigarettes. Prenatal THC exposure via e-cigarettes significantly delayed the age for offspring to achieve a successful trial, a developmental milestone in rats. Although offspring with prenatal THC exposure showed this delay in the first couple of days of testing (PD 12-13), it is important to note that they did eventually catch up to the performance of other groups in this task (PD 15). However, these same offspring showed impairments on a motor task that requires coordination and balance later in life, from PD 30-32 (a period similar to human adolescence). Offspring exposed to prenatal THC via e-cigarettes required significantly more trials to successfully traverse the parallel bars and had a lower success ratio on the first day of the task, achieving lower maximum widths between bars. These impairments in early motor development and adolescent motor coordination following prenatal THC exposure were not exacerbated by alcohol, nor were they dependent on sex, suggesting an overall motor impairment induced by prenatal exposure to THC.

These results differ from our report that combined ethanol and cannabinoid (CP-55,940) during the 3^rd^ trimester advances motor development, with long-term motor impairments in females. In addition to the differing cannabinoids (THC vs CP-55,940) and administrative levels and routes, these studies also varied in developmental timing of exposure, which would impact different aspects of cannabinoid receptor development in several motor-related brain areas. For example, cannabinoid type 1 (CB1) receptors play roles in embryonal implantation, neuronal development, and early synaptic communication as early as GD 11-14 in rodents (Berghuis et al., 2007; Harkany et al., 2007); during this time, prenatal cannabis exposure may impair early developmental processes, including motor function. In the present study, exposure occurred between GD 5-20, consistent with clinical and preclinical studies reporting that prenatal cannabinoid exposure is associated with decreased early motility (Fried, 1976), impaired early motor skills (Richardson et al., 1995), and impaired motor coordination (Shabani et al., 2011). In contrast, CB1 binding in the GABAergic neurons of rodents double rapidly postnatally, from PD 0-14, in the cerebellum (Diana et al., 2002), basal ganglia, and cortex (Berrendero et al., 1999), which are important for motor control; during this time, CB1 receptor activation may up-regulate GABA activity and enhance performance (Benagiano et al., 2007; Chiu et al., 2005). This is consistent with the advanced sensorimotor development we found with administration with a synthetic cannabinoid during the human third trimester equivalent (PD 4-9) (Breit et al., 2019a). Similar motor advancement has been seen in clinical studies (Fried & Watkinson, 1990), although the exact timing of exposure cannot always be confirmed in clinical retrospective studies. Nevertheless, even when sensorimotor function was advanced early in development, long-term motor function was not affected or was impaired. Importantly, fetal development requires a careful timing of events, and any changes in neuronal development can lead to later-life consequences (Wolosker et al., 2008). Thus, the overall conclusion is that prenatal cannabis exposure may alter motor development and coordination among offspring, and those alterations may vary based on the timing of administration.

In contrast, distinct interactive effects of prenatal alcohol and THC exposure via e-cigarettes were seen in the open-field activity chambers. Both clinical and preclinical data examining the effects of cannabinoid exposure during early development on activity levels are mixed, with data in each field supporting increased activity (Borgen et al., 1973; Breit et al., 2019b; Fried et al., 1992; Goldschmidt et al., 2000; Mereu et al., 2003; Navarro et al., 1994; Rubio et al., 1995) and no change in activity levels (Brake et al., 1987; Fried & Watkinson, 1988; Hutchings et al., 1989; Vardaris et al., 1976). Prenatal alcohol exposure has been linked to greater levels of attention deficit hyperactivity disorder symptoms (Nash et al., 2006; O’malley & Nanson, 2002; Popova et al., 2016) and increased hyperactivity in rodents (Bond, 1981; Breit et al., 2019b; Ryan et al., 2008; Thomas et al., 2007), similar to clinical findings among individuals with FASD (Dörrie et al., 2014; Mattson et al., 2011). The current study found modest increases in locomotor activity following either prenatal alcohol or THC among females. However, only the combination of prenatal alcohol and THC increased activity levels only in males. This result is similar to our previous work showing that early developmental exposure to either alcohol or a synthetic cannabinoid (CP-55,940) during the 3^rd^ trimester equivalent brain growth spurt increased adolescent activity levels, but that the combination exacerbated hyperactivity among offspring, by reducing habituation in locomotor activity during the testing session (Breit et al., 2019b). Importantly, in both studies, co-exposure to alcohol and cannabinoids increased blood alcohol levels more than the alcohol dose alone in both pregnant dams (Breit et al., 2020) and neonates (Breit et al., 2019b). Given that the severity of fetal alcohol spectrum disorders may be dose-dependent (Jacobson et al., 1998; Kelly et al., 1987; Maier & West, 2001), it is possible that the interactive effects of alcohol and cannabinoid exposure on activity levels could be related to higher blood alcohol concentrations. Research examining the effects of combined prenatal alcohol and cannabis exposure on long-term behavioral development among offspring is incredibly sparse, and this possibility should be further examined using dose-dependent studies.

Not only did combined prenatal exposure to alcohol and THC increase overall activity levels, but interactive effects were also seen in behaviors specifically in the center of the open-field chambers. Combined exposure to alcohol and THC via e-cigarettes during gestation increased the number of entries into the center, time spent in the center, and the distance traveled in the center of the chambers among male offspring. Rodents are naturally aversive to open spaces (such as the chamber center), and thus increased center-related activities could either be a by-product of overall increased locomotor activity, but could also be related to alterations in anxiety-related behaviors such as decreased risk assessment or increased impulsivity (Prut & Belzung, 2003). In the current study, it is unclear why this interactive effect was only observed among male offspring, given that there is very little known about the effects of combined prenatal alcohol and cannabis exposure in general. However, some clinical (Ouellet-Morin et al., 2011) and preclinical data (Hellemans et al., 2010) do suggest that males are more susceptible to these types of alterations in activity levels and anxiety-related domains following prenatal alcohol exposure comparted to females.

One limitation of this study was that it utilized single doses of both alcohol and THC (Breit et al., 2020). It will be important to understand how prenatal alcohol and THC at wider ranges, including the high doses of THC currently consumed, influence behavioral development. Although these doses are not as high as those published in substance abuse studies, effects of moderate doses are important to research since they represent what is more commonly used among women of child-bearing age. In addition, this study focused on prenatal THC exposure, the primary psychoactive component of cannabis; importantly, there are more than 500 chemical compounds, including over 100 naturally-occurring cannabinoids (Radwan et al., 2017). Thus, we acknowledge the outcomes of this study may not generalize to all types of prenatal cannabis exposure. Lastly, although e-cigarettes are a popular route of administration for THC, including among pregnant women, we recognize that vapor inhalation is not as clinically relevant of an administration route for alcohol. We chose to expose pregnant dams to alcohol via vapor inhalation since they would already be exposed to vapor via e-cigarettes, and we wanted to minimize additional stress. Importantly, we found similar effects using this alcohol vapor inhalation paradigm (Breit et al., 2020) on maternal weight gain and offspring eye opening as we have with maternal intragastric intubation (Thomas et al., 2009). In addition, we have also found similar pharmacokinetic effects with combined alcohol and synthetic cannabinoid exposure via neonatal alcohol intubations and cannabinoid intraperitoneal injections (Breit et al., 2019a).

### 4.1.1. Conclusions

In summary, these results suggest that prenatal exposure to alcohol and THC via e-cigarettes have domain-specific effects on motor-related behaviors. Prenatal THC exposure decreased offspring body weights throughout early and adolescent development, regardless of sex. In addition, prenatal THC exposure delayed both early sensorimotor development and adolescent motor coordination. Prenatal alcohol exposure at this dose did not alter body growth, motor development, or motor coordination, and produced modest effects on activity level. In contrast, combined prenatal exposure to alcohol and THC increased activity levels. Taken together, prenatal cannabis exposure may lead to impair motor performance, whereas combined exposure with alcohol may lead to hyperactivity and altered impulsivity. These results have important implications for public health and policy, especially since alcohol and cannabis are often consumed together, and nearly 40% of pregnancies in the United States are unplanned (Mosher et al., 2012).

## Acknowledgments

This work was supported by a NIAAA grant AA025425 to Dr. Jennifer D. Thomas, a NIAAA training grant T32AA007456-38 to Dr. Kristen R. Breit, and a NIAAA Loan Repayment Program award to Dr. Kristen R. Breit. Special thanks to Maury Cole at La Jolla Alcohol Research, Inc. (San Diego, CA) for building the vapor inhalation equipment and Dr. Michael Taffe for vapor inhalation advisement. Thank you to the NIDA Drug Supply for providing all THC used in this study. Lastly, we want to recognize members of the Center for Behavioral Teratology at San Diego State University for assisting in data collection and interpretation, particularly the instrumental efforts of Brandonn Zamudio, Ivette Gonzalez, Bahar Sabouri, and Valery Quinonez.

## Author Credits

Dr. Kristen R. Breit: Conceptualization, Formal analysis, Investigation, Methodology, Project Administration, Visualization, Roles/Writing – original draft; Cristina Rodriguez: Investigation, Methodology, Project administration, Validation; Annie Lei: Investigation, Methodology, Validation; Samirah Hussain: Investigation, Methodology, Validation; Dr. Jennifer D. Thomas: Conceptualization, Funding acquisition, Supervision, Writing – review & editing

